# Resilient functioning is associated with altered structural brain network topology in adolescents exposed to childhood adversity

**DOI:** 10.1101/2023.05.05.538901

**Authors:** Nadia González-García, Elizabeth E.L. Buimer, Laura Moreno-López, Samantha N. Sallie, František Váša, Sol Lim, Rafael Romero-Garcia, Maximilian Scheuplein, Kirstie Whitaker, Peter B. Jones, Ray Dolan, NSPN consortium, Peter Fonagy, Ian Goodyer, Ed Bullmore, Anne-Laura van Harmelen

**Author notes:** shared first authorship (Nadia González-García, Elizabeth E.L. Buimer). + NSPN author list: https://nspn.org.uk/nspn-team/. Corresponding Author: Prof. Anne-Laura van Harmelen Pieter de la Court Building (room 4B48) Wassenaarseweg 52 2333 AK Leiden, Netherlands. **Author Contributions:** LML and NGG analysed the imaging data and drafted the manuscript with EELB, SNS, MS and ALvH. ALvH conceptualized and designed the study, analysed the behavioural data, and reviewed and revised the manuscript. All authors approved the final manuscript and agreed to be accountable for all aspects of the work presented. **Competing Interest Statement:** ETB is a member of the scientific advisory board for Sosei Heptares and a consultant for GlaxoSmithKline.

## Abstract

Childhood adversity is one of the strongest predictors of adolescent mental illness. Therefore, it is critical that the mechanisms that aid resilient functioning in individuals exposed to childhood adversity are better understood. Here, we examined whether resilient functioning was related to structural brain network topology. We quantified resilient functioning at the individual level as psychosocial functioning adjusted for the severity of childhood adversity in a large sample of adolescents (N=2406, aged 14-24). Next, we examined nodal degree (the number of connections that brain regions have in a network) using brain-wide cortical thickness measures in a representative subset (N=275) using a sliding window approach. We found that higher resilient functioning was associated with lower nodal degree of multiple regions including the dorsolateral prefrontal cortex, the medial prefrontal cortex, and the posterior superior temporal sulcus (*z* > 1.645). During adolescence, decreases in nodal degree are thought to reflect a normative developmental process that is part of the extensive remodelling of structural brain network topology. Prior findings in this sample showed that decreased nodal degree was associated with age, as such our findings of negative associations between nodal degree and resilient functioning may therefore potentially resemble a more mature structural network configuration in individuals with higher resilient functioning.

## Introduction

Childhood adversity (CA) refers to a range of negative experiences throughout childhood and adolescence such as parental psychopathology, peer victimization, and various forms of parental maltreatment (e.g., neglect or overt maltreatment). CA experiences are one of the strongest predictors of mental health problems (Green et al., 2010), possibly through their impact on the developing brain. According to the social transactional model of psychiatric vulnerability (McCrory et al., 2022), CA experiences shape fronto-limbic development and related socio-emotional functioning to aid survival in high threat environments. For instance, in the context of an abusive home-environment it may be adaptive for a child to rapidly detect when a parent is angry, to expect negative feedback, and to adjust their behaviour and emotions accordingly. However, in non-threatening social environments such socio-emotional functioning adaptations may inadvertently evoke social problems and generate social stress, and ultimately lead to mental health problems in later life (McCrory et al., 2022).

Fortunately, not all individuals who have experienced CA develop mental illness; rather, a substantial proportion of individuals exposed to CA function resiliently later in life. Resilience refers to the capacity of a system (e.g., a brain, a child, a family, a community) to successfully adapt to challenges that threaten the function, survival, or development of that system (Masten et al., 2021; Masten & Monn, 2015). Resilience in the context of CA, when the stressor has already taken place, refers to an outcome of positive mental health *functioning* on a given trajectory or at a given point in time. Such resilient functioning should be assessed across mental health domains, given the non-specific negative impact of CA (Masten & Monn, 2015), and should be better than others with similar severity of CA experiences (van Harmelen et al., 2017). Resilient functioning in individuals exposed to CA is aided by an array of separate yet interrelated protective social and cognitive influences such as positive parenting, and social support, high self-esteem, low rumination (Fritz et al., 2018; Kalisch et al., 2017, 2019; van Harmelen et al., 2017; 2020). Furthermore, it is thought that these socio-cognitive protective factors interact with brain structure and functioning (Ioannidis et al., 2020). Recent reviews of the literature suggest that resilient functioning may be facilitated by larger hippocampal structure and increased functional connectivity between limbic regions and the central executive network (Moreno-Lopez et al., 2020). Although these studies provide important insights about the neurobiology that may aid resilient functioning, recent advances in neuroscience indicate that cognitive and emotional processes are not merely facilitated by specific regions but emerge through the interaction of brain networks (Krendl & Betzel, 2022).

Brain networks can be constructed from structural or functional neuroimaging data (Krendl & Betzel, 2022). Structural covariance networks are thought to overlap with functional networks (Zielinski et al., 2010), and can be examined using graph theoretical approaches, such as structural covariance (Bullmore & Bassett, 2011; Kaiser, 2011; Rubinov & Sporns, 2010). Structural covariance reflects the inter-individual (Alexander-Bloch et al., 2013; Vijayakumar et al., 2021) or intra-individual (Seidlitz et al., 2018; Yun et al., 2016) covariation in brain morphology (for example cortical thickness) between different regions (nodes). Importantly, inter-individual structural covariance may reflect co-ordinated brain development (Alexander-Bloch et al., 2013; Khundrakpam et al., 2013; Vijayakumar et al., 2021). During puberty and adolescence, the cerebral cortex becomes thinner (Wierenga et al., 2014) and white matter tracts become more densely myelinated (Miller et al., 2012) suggesting a progressive refinement of neural connections through ongoing neural regressive events such as pruning (Kaiser et al., 2011; Zielinski et al., 2010; Miller et al., 2012; Alexander-Bloch et al., 2013). The transition from childhood to adolescence is characterized by global increases in structural covariance of cortical thickness followed by reductions into mid-adolescence (Vijayakumar et al., 2018). Structural covariance continues to decrease through late adolescence before plateauing in the early twenties, which corresponds to the prolonged maturation of association cortices (Váša et al., 2018). The associated developmental changes in structural covariance during later adolescence, such as cortical thinning, have been related to reductions in nodal degree, the number of connections that brain regions have in a network (Váša et al., 2018). Such development is thought to be shaped by genetic as well as environmental influences (Whitaker et al., 2016).

Reviews on the neurobiology of resilience show an overall lack of consistent findings which could be the result of different conceptualizations of resilience (Eaton et al., 2022; Zhang et al., 2023; Leal & Silvers, 2020). In general, reviews point to neural circuits involved in emotion regulation and reward (Eaton et al., 2022; Leal & Silvers, 2020), fronto-subcortical networks (Zhang et al., 2023) and the emotional brain (Moreno-López et al., 2020). To date, only a few studies have investigated the relationship between resilience in individuals exposed to CA and structural brain networks (reviewed in Moreno-López et al., 2020). Ohashi and colleagues (2019) found reduced nodal efficiency in resilient individuals in the amygdala and 8 other nodes compared to susceptible individuals exposed to CA using diffusion tensor imaging and tractography. Comparing groups of non-maltreated youth, maltreated youth with PTSD and maltreated youth without PTSD, Sun et al., (2019) found larger centrality (importance of a region within a network) in the right frontal pole in maltreated youth without PTSD symptomatology compared to non-maltreated youth and maltreated youth with PTSD based on a structural covariance derived from cortical thickness estimates. The frontal pole plays a role in adapting and updating reward processing models in response to the environment (Kovach et al., 2012). In a study with a similar design, maltreated youth without PTSD (versus with PTSD) showed larger centrality in right orbitofrontal cortex (Sun et al., 2018), a region critical for evaluation, affect regulation and reward-based decision-making (Fettes et al., 2017). Thus, resilient functioning in individuals exposed to CA is likely related to altered structural covariance patterns. However, these studies estimated resilience at the group-level, and were not able to relate the findings to individual level of resilient functioning, limiting the generalizability of these findings.

Hence, little is known about the structural network topology related to the level of resilient functioning in individuals exposed to CA, particularly in young people when the brain is in development. In doing so, appropriate quantification of resilient functioning after CA must keep in mind the following aspects. Firstly, given the negative impact of CA on a range of mental health and wellbeing outcomes, it is important that resilient functioning incorporates functioning across these psychological and social (‘psychosocial’) domains of functioning (Masten & Monn, 2015). Such resilient functioning across domains should further take into account what someone has experienced, as individuals with similar psychosocial functioning may differ in their degree of resilient functioning when one has experienced more severe CA than the other. Finally, as CA is a highly clustered experience, where different types of adversity often co-occur, it is important to take all CA experiences into account when examining CA. To do so, we build on previous work (Ioannidis et al., 2020; van Harmelen et al., 2017, 2021) and use data reduction techniques (principal component analyses) to derive a single estimate for psychosocial functioning that summarises low to high functioning across multiple measurements, and use the same approach to calculate a single estimate that summarises the severity of all experiences of childhood family adversity (CFA) in a community cohort of healthy young people with low to moderate CFA (N=2406, aged 14-24). Next, we regressed the estimate for CA onto the estimate for psychosocial functioning. In doing so, individual level resilient functioning can then be inferred from the residuals of the relation between CA and psychosocial functioning - the extent to which an individual is functioning better than expected given their CA experiences (implying resilient functioning, green lines) or worse than expected (implying vulnerable functioning, red lines) (Fig. 1b, see (Ioannidis et al., 2020; van Harmelen et al., 2017, 2021). The aim of this study is to examine whether such resilient functioning in young people exposed to CFA is associated with altered structural network topology. To do so, we use a sliding window method (See Figure 1c); a novel approach so far only used to estimate the structural network topology of neurodevelopmental trajectories (Alexander-Bloch et al., 2013; Ohashi et al., 2019). Here, we use the sliding window method to be able to use resilient functioning as a continuous measure and test for robustness of its association with nodal degree by repeating the structural covariance analyses across different subsamples. By altering the window width and step size with each iteration, our findings get independent of the parameters used for the sliding window method. Given the importance of socio-emotional functioning in mental health vulnerability in adolescents and adults exposed to CA (McCrory et al., 2022; Moreno-López et al., 2020), we hypothesized that higher level of resilient functioning would be associated with changes in nodal degree of cortical brain regions that help guide socio-emotional functioning. Building on previous work in this sample (Váša et al., 2018; Whitaker et al., 2016), we chose to investigate nodal degree in structural covariance networks derived from cortical thickness estimates since this measure was shown to decrease with age in adolescence, and this decrease was associated with corresponding myelination changes in the association cortices (Váša et al., 2018). Furthermore, we focused on CFA rather than the broader CA as our available data contains questionnaires that focus on the family environment. Data on other experiences of CA, such as bullying, racism or poverty, were not included here.

**Figure 1.**
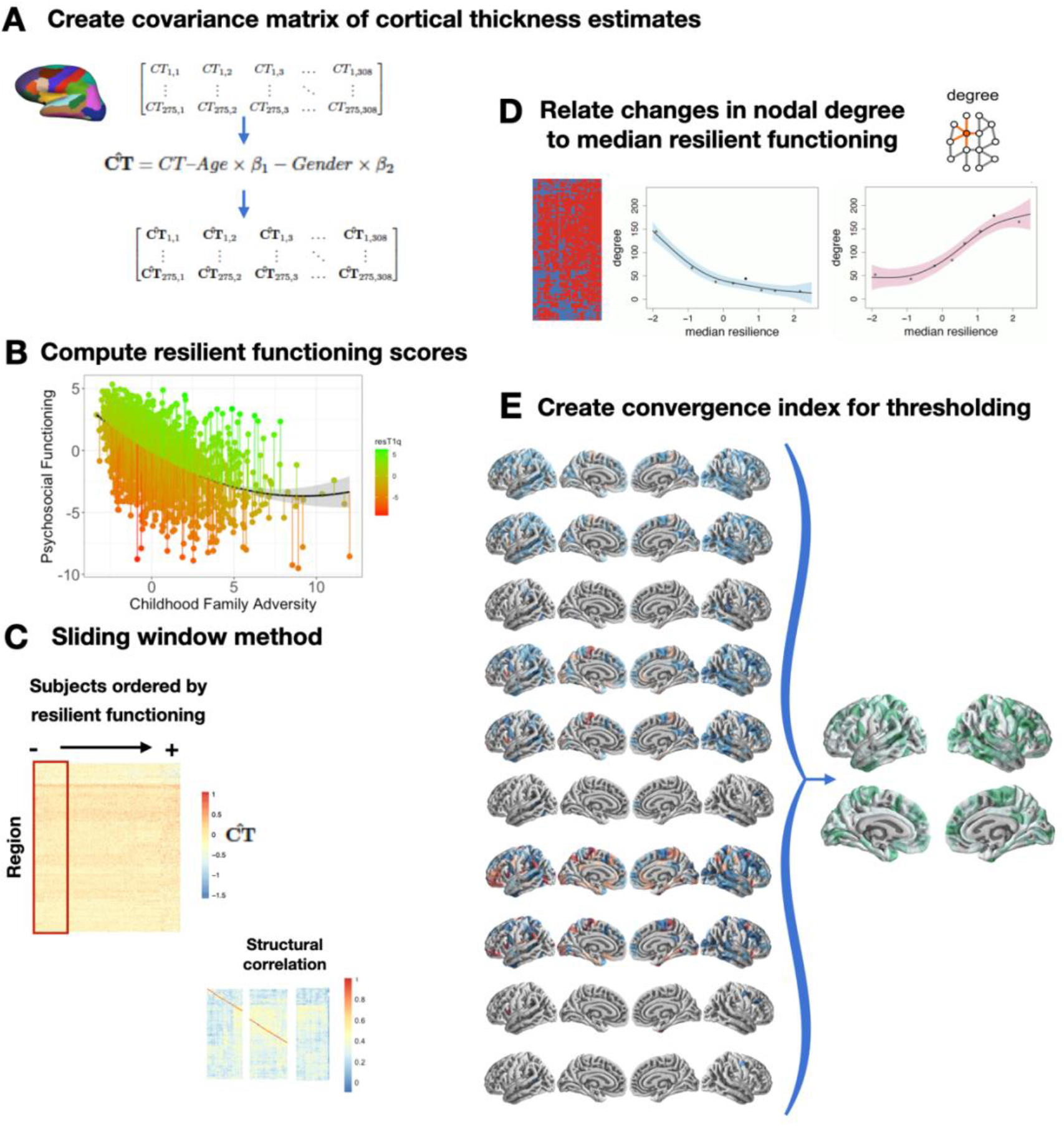
Study design. A) A covariance matrix of the cortical thickness (CT) measures for 308 parcellations in 275 participants is created. Next, the data matrix is substituted by the residuals of the linear regression to remove variation related to age, gender, and intra-cranial volume. B) Next, resilient functioning scores are created based on the NSPN sample (N = 2406). The figure shows the extent to which an individual functioned better than expected (‘high, or resilient’; green lines), or worse than expected (‘low or risk’ red lines), than others with similar childhood family adversity experiences. Note that higher residual scores reflect more resilient functioning, and that both X and Y axes represent factor scores with Mean=0 and SD=1. C) Next, a sliding window method was applied with varying window sizes (red box), to assess how resilient functioning was related to changes in the nodal degree of the network overlapping structural networks of 275 participants. CT values of each region were cross correlated with windows containing the same numbers, we used bootstrapping to threshold the network. D) Then, we evaluated linear regional changes in node degree as a function of the median resilient functioning. E) We varied these parameters to explore consistency in the results using a “Convergence Index”, considering all nine combinations of window widths (40, 60, 80) and step sizes (5, 10, 20), plus one further combination of ww = 60 and ss = 30. Convergence indices were calculated for each of the brain parcellations, where the index represents the number of times the region is associated with resilient functioning for each of the above combinations. Thus, a convergence index of 10 indicates that the region was associated with resilient functioning every run and a convergence index of 0 indicates that it was never associated.

## Methods

### Study design and participants

Participants were part of the Neuroscience in Psychiatry Network (NSPN) study: a multi-centre accelerated longitudinal community cohort study focusing on normative adolescent to young adult development between the ages 14-24. The NSPN cohort (*N* = 2406) was recruited from schools, colleges, National Health Service (NHS) primary care services, and direct advertisement in north London and Cambridgeshire. Maintaining the same gender and ethnicity balance as in the main sample, 301 participants were invited for an MRI scan (Whitaker et al., 2016). For this manuscript, we excluded those individuals with a lifetime history of brain damage, epilepsy, genetic syndromes, and premature birth (*N* = 26), leaving a total sample of 275 individuals for the analyses of brain imaging data below.

The inclusion criteria for the MRI subset were that the participants should be aged between 14 and 24 years; able to understand written and spoken English; have normal or corrected-to-normal vision; and able to give informed consent for participation in the study. The exclusion criteria were current treatment for psychiatric disorders, drug dependence, alcohol dependence, current or previous neurological disorders, brain trauma including epilepsy, head injury causing loss of consciousness, learning disability requiring specialist educational support and/or medical treatment and standard MRI contraindications. Individuals with previous or lifelong psychiatric disorders were not excluded, except if they were in current treatment for these disorders. The study was approved by the Cambridgeshire and Peterborough Foundation Trust and University of Cambridge research ethics committees (REF 12/EE/0250). All participants (and their caregivers) were briefed about the study aims and protocols and signed an informed consent form.

### Acquired data

To assess resilient functioning, we relied on data from questionnaires on psychological functioning, CFA, socio-demographic status, family and educational or occupational environments, and sub-clinical psychopathology in the NSPN sample (*N* = 2406). Assessments for the MRI sub study (*N* = 275) further included a day of clinical, cognitive, and MRI assessments at the University of Cambridge or University College London sites.

#### Measures of psychosocial functioning

Psychosocial functioning was based on *all* measures included in the NSPN home questionnaire pack that assessed any aspect of psychological and social functioning. This included measures of psychiatric symptomatology, personality traits, and mental wellbeing. Below we provide an overview of the specific measured used, please refer to van Harmelen et al., (2017) and Supplementary material for a description of the measures. Psychosocial functioning was assessed with sum scores from the Mood and Feelings Questionnaire (MFQ; (Angold et al., 1996)), Revised Children’s Manifest Anxiety Scale (RCMAS; (Reynolds & Richmond, 1997)), Short Leyton Obsessional Inventory (S-LOI;(Bamber et al., 2002)), Child Behavior Checklist (CBCL; (Achenbach, 1991)) and Kessler Psychological Distress Scale (K10; (Kessler et al., 2002)), the Adolescent Psychopathy Screening Device (APSD; (Frick et al., 2000)), Child and Adolescent Dispositions Scale (CADS; (Lahey et al., 2008)), the Inventory of Callous-Unemotional Traits (ICU; (Roose et al., 2010), Schizotypal Personality Questionnaire (SPQ; (Raine, 1991)), and the Barratt Impulsiveness Scale (BIS-11; (Stanford et al., 2009)), and the Warwick-Edinburgh Mental Well-being Scale (WEMWBS; (Tennant et al., 2007)).

#### Measures of childhood family adversity

Childhood Family Adversity (CFA) scores included *all* measures in the NSPN home questionnaire pack that assessed any aspect the home environment whilst growing up. As such, CFA within the family environment was assessed with the Alabama Parenting Questionnaire (APQ; (Elgar et al., 2007)) and the Measure of Parental Style (MOPS; (Parker et al., 1997)). The types of experiences assessed with the APQ and MOPS include parental abuse and neglect, and more general parenting behaviours (i.e., positive parenting, inconsistence, indifference, and control). Please refer to van Harmelen et al., (2017) and Supplementary material for a description of the measures.

#### MRI acquisition

Participants underwent structural MRI (3T) using the multi-parameter mapping sequence (Weiskopf et al., 2013) in Cambridge (2 sites) or London. All sites used identical scanners (Siemens Magnetom Tim Trio), sequences, and protocols. The setup, acquisition and post-processing have been previously described (Weiskopf et al., 2013). Briefly, the multi-parameter mapping (MPM) protocol includes three multi-echo 3D FLASH (fast low angle shot) scans, one radiofrequency (RF) transmit field map and one static magnetic (B0) field map scan. Multiple gradient echoes were acquired with alternating readout polarity at six equidistant echo times (TE) between 2.2 and 14.7 ms for both acquisitions. Other acquisition parameters were: 1 mm isotropic resolution, 176 sagittal partitions, field of view (FOV) = 256×240 mm, matrix = 256 x240×176, parallel imaging using GRAPPA factor 2 in phase-encoding (PE) direction (AP), 6/8 partial Fourier in partition direction, non-selective RF excitation, readout bandwidth BW = 425 Hz/pixel, RF spoiling phase increment = 50Å. The total acquisition time was ∼25 min. Participants were instructed to lie still and rest during the scan.

#### MRI processing

MR images were processed using the Freesurfer pipeline (v5.3.0), including skull-stripping, and segmentation of cortical grey and white matter (Fischl et al., 2002). After quality control, three participants were excluded from further analysis because of movement artifacts, which prevented accurate surface reconstructions and reconstruction of the cortical surface and grey-white matter boundary (for more detail see Whitaker et al., (2016)). Parcellation of cortical gray matter regions was based on the anatomical borders of 308 equally sized regions (159 in each hemisphere) of 500mm^2^ that were constrained by the anatomical boundaries according to the Desikan-Killiany atlas (Desikan et al., 2006). We opted to use this particular atlas because it comprises an optimal spatial scale for graph theory analysis (Romero-Garcia et al., 2012). Average CT was extracted for each of the 308 regions in each participant.

### Analyses

#### Missing data handling

All analyses were conducted using R version 3.4.1 ʽSingle Candle’ of the Lavaan package (Rosseel, 2012). From the NSPN cohort (*N* = 2406), all behavioural measures were complete for 1907 participants. The subset with no missing data did not differ from the larger sample in terms of age (*t*(4091) = -0.027, *p* = 0.97), gender (*χ*^2^ = 0.01, df = 1, *p* = 0.92), socio economic status (SES; index of multiple deprivation based on participant postcodes; *t*(4038) = -1.29, *p* = 0.19), or ethnicity (*χ*^2*2*^ = 3.65, df = 5, *p* = 0.60). Thus, missing data on the measures used in the below analyses was imputed using the Amelia package in R (Honaker et al., 2011). We calculated resilient functioning scores (as per the below description) within all 5 imputed datasets. These resilient functioning scores were highly correlated (*r*’s > 0.9, see supplemental Table S1 for specifics). Therefore, for this manuscript, we used the resilient functioning scores that were calculated using data from the first imputation sample.

#### Resilient functioning scores

Following the procedure detailed in van Harmelen et al., (2017), we estimated resilient functioning using the ‘residual method’ on the imputed dataset for the NSPN cohort (N = 2406). Please refer to Ioannidis et al. (2020) for a detailed discussion of the benefits and drawbacks of this approach to quantify resilient functioning, and Cahill et al., (2022) for external validation of this approach which has shown good psychometric properties. Using this method we previously showed that adolescent friendships predict resilient functioning in two large independent samples; the N=2406 NSPN sample (van Harmelen et al., 2017), and in the N=1238 Roots sample (van Harmelen et al., 2021). We used principal component analysis (PCA) to computed individual psychosocial functioning scores using standard-normally transformed individual total scores across a range of measures (see Table 1; MFQ, RCMAS, S-LOI, K10, CBCL, APSD, CADS, ICU, SPQ, BIS-11, and WEMWBS). We also utilized PCA to calculate individual levels of CFA severity using standard-normally transformed sum scores for the MOPS and the APQ sub-scales (see Table 1) within the entire NSPN sample (N = 2409). Resulting CFA factor scores were regressed onto the psychosocial functioning factor scores, and the best fitting regression model (in this case, quadratic) was obtained. The residuals from this model reflect how much better or worse individuals are functioning when compared to others with similar CFA scores. As such, these residual scores can be interpreted as a proxy to indicate individual degree of ‘vulnerable to resilient’ functioning (from here, for brevity we refer to this as ‘resilient functioning’), with higher scores reflecting better psychosocial functioning relative to the level of CFA. Next, individual resilient functioning scores were extracted for the MRI cohort (N=275) and utilized in subsequent analyses.

**Table 1.**
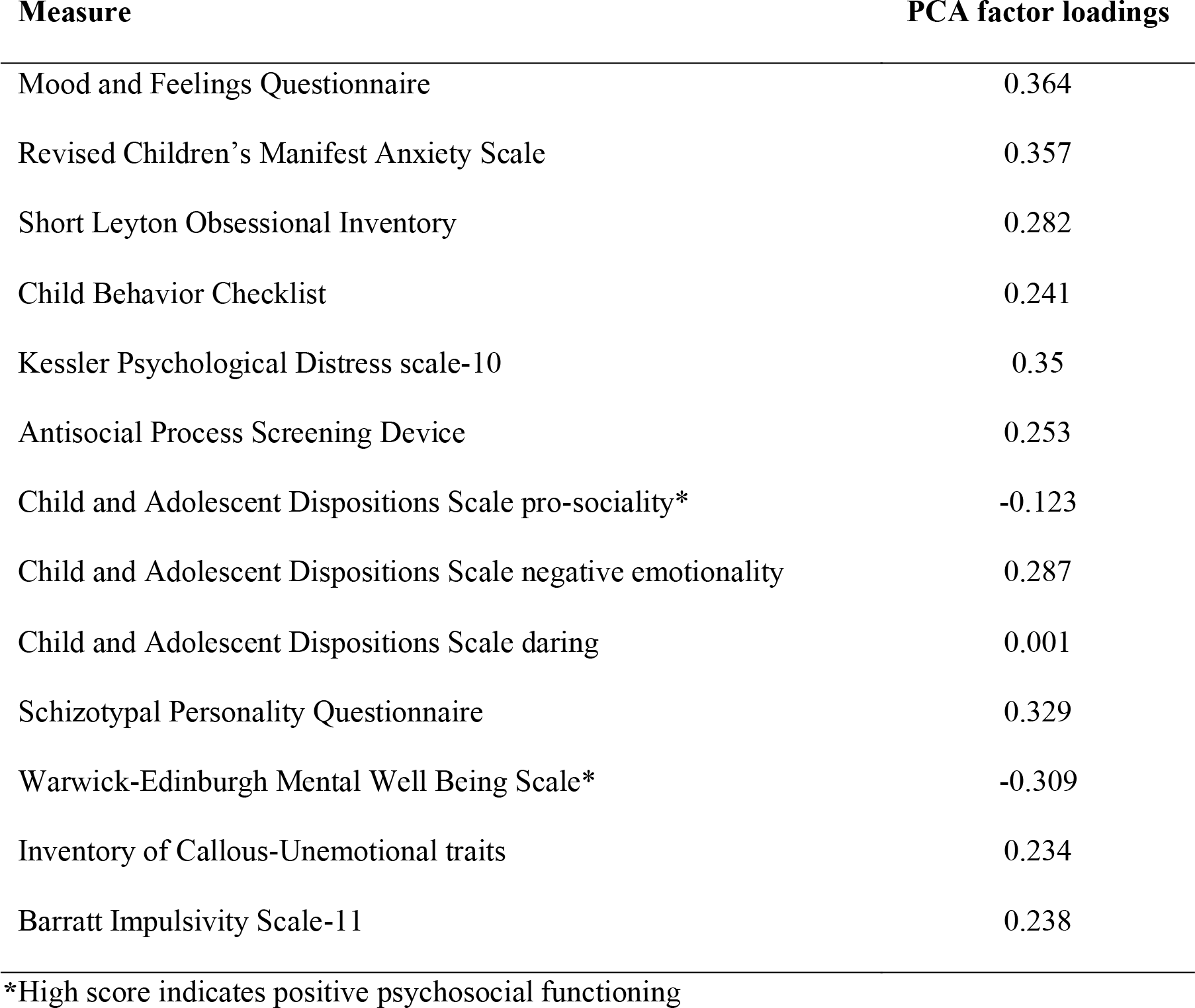
Factor loadings for the PCA’s for psychosocial functioning.

| Measure | PCA factor loadings |
| --- | --- |
| Mood and Feelings Questionnaire | 0.364 |
| Revised Children's Manifest Anxiety Scale | 0.357 |
| Short Leyton Obsessional Inventory | 0.282 |
| Child Behavior Checklist | 0.241 |
| Kessler Psychological Distress scale-10 | 0.35 |
| Antisocial Process Screening Device | 0.253 |
| Child and Adolescent Dispositions Scale pro-sociality* | -0.123 |
| Child and Adolescent Dispositions Scale negative emotionality | 0.287 |
| Child and Adolescent Dispositions Scale daring | 0.001 |
| Schizotypal Personality Questionnaire | 0.329 |
| Warwick-Edinburgh Mental Well Being Scale* | -0.309 |
| Inventory of Callous-Unemotional traits | 0.234 |
| Barratt Impulsivity Scale-11 | 0.238 |

\*High score indicates positive psychosocial functioning

#### Structural covariance

To estimate each structural network, we used Pearson’s correlation coefficients on the cortical thickness (CT) values estimated from Freesurfer based on structural MRI. Cortical thickness estimates were corrected for intracranial volume. Further, we performed a linear regression on regional CT values to remove effects of age and gender on cortical thickness. The residuals of this regression then replaced the raw values in the CT data matrix. These detrending steps were implemented to remove potentially confounding inter-individual variation related to age and gender and intracranial volume. This method has been utilized by others (Melie-Garcia et al., 2018). Next, we used bootstrapping to threshold the network. Using this method, and for each window, an equal number of participants were resampled with replacement to construct 1000 bootstrapped structural networks. We then examined whether there were significant relations between each pair of regions across all bootstrapped networks. Consistent relationships between a pair of regions (at *p* < .001 adjusted for the False Discovery Rate (FDR) at the pair level (Váša et al., 2018)) were retained and the remaining relationships were discarded. For comparison, the main analyses were repeated without the detrending step for age and gender (Supplement).

#### Sliding window method

We applied a sliding window method to assess how resilient functioning was related to changes in the nodal degree of the network, defined as the number of edges connected to a node (Figure 1). CT values of each region were cross correlated with windows containing the same numbers of participants and moved across resilient functioning scores by stepwise increases. At each step (within each window) we estimated a structural covariance network. For more information on the sliding window technique see Váša et al. (2018). The selection of sliding window parameters, including window width (*ww*) and step size (*ss*) in units of number of participants involved several trade-offs. The number of windows was defined using the following equation:

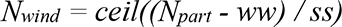

Where *N_part_* was the number of participants (*n* = 275), and *ceil* the ceiling function, which rounds non-integer fractions to the smallest integer larger than said fraction. The *ss* and *ww* define the number of windows and this in turn has an impact on the sensitivity of the analyses. Therefore, we varied these parameters to explore consistency in the results, considering all nine combinations of window widths (40, 60, 80) and step sizes (5, 10, 20), plus one further combination of *ww* = 60 and *ss* = 30, overlapping structural networks of 275 participants. We assessed the network topology changes by measuring the nodal degree. We then evaluated linear regional changes in nodal degree as a function of the median resilient functioning with Akaikes information criterion (AIC) and corrected for multiple comparisons (adjusted for the false discovery rate, *p* < .05). We ran permutation tests at the level of the windows without reconstructing the correlation matrices to see if the regional effects of change in structural correlation as a function of resilient functioning was valid. A convergence index was calculated for each of the brain region under consideration, where the index represents the number of times the region is associated with resilient functioning for each of the above combinations. Thus, the convergence index of a cortical region 10 indicates that the region is always associated with resilient functioning and the convergence value of 0 of another region (or node) indicates that it never is (see Figure 1). Next, we conducted a standardization to define the significant regions associated with resilient functioning (*z* > 1.645).

### Availability of data and code

Data and code for the analyses will be available upon reasonable request at the University of Cambridge repository (https://www.repository.cam.ac.uk/). We uploaded the main results in the neuroimaging repository neurovault.org (Gorgolewski et al., 2015). The 1_6CI.nii.gz upload (https://identifiers.org/neurovault.image:785762) corresponds to the mean of the ß-estimates after thresholding of the main effect of resilient functioning on local nodal degree (Figure 2).

**Figure 2.**
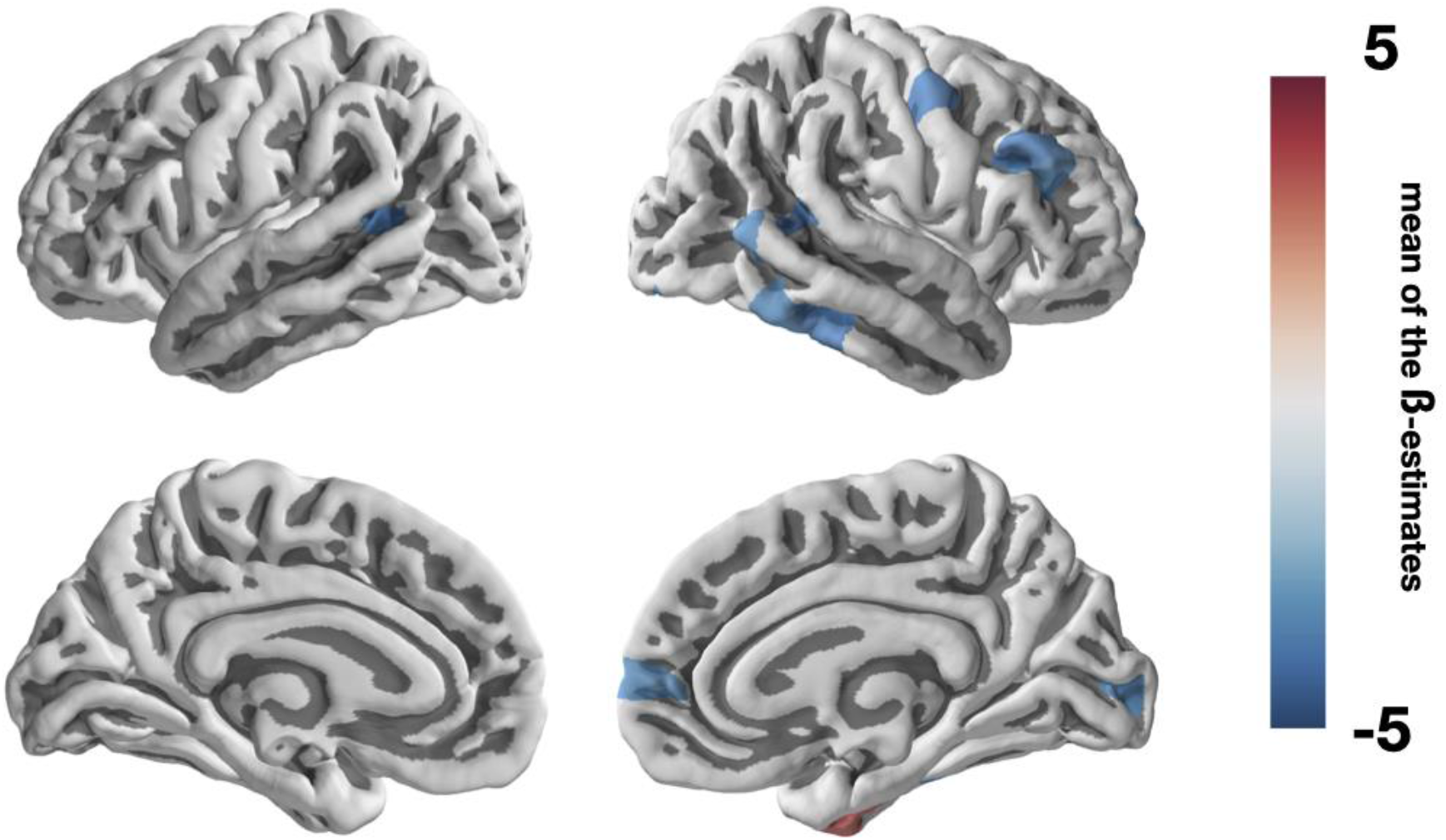
Brain parcellations with positive or negative correlations between nodal degree and resilient functioning. Cortical regions where resilient functioning was significantly convergently associated with nodal degree after thresholding based on the convergence index (z > 1.645). The colorbar represents the mean of the β-estimates with positive correlations in red and negative correlations in blue.

## Results

### Principal component scores

The first principal component of PCA for psychosocial functioning explained 44% variance across all psychological functioning measures (SD = 2.41, see Table 1 for all factor loadings). A higher score on the first component score was related with poorer psychosocial functioning, therefore, individual scores were subsequently inverted so that higher scores would indicate better psychosocial functioning.

The PCA for CFA resulted in a first component score that explained 37% variance (SD = 2.02, see Table 2 for specifics). Here, higher scores were related to lower CA and were subsequently inverted to indicate more CFA. As the score for CFA included a few positive parenting scales, we repeated the PCA without the positive parenting subscale. After removal of this subscale the explained variance of our principal component was reduced with 0.3% (from 37.2% to 36.9%). Furthermore, principal component scores with and without this subscale correlated highly (*r* = .98, *t* = -331.5, df = 2404, *p*-value < 2.2e-16). Therefore, we decided to leave the positive parenting subscales in the PCA, in line with previous work (van Harmelen et al., 2017).

**Table 2.**
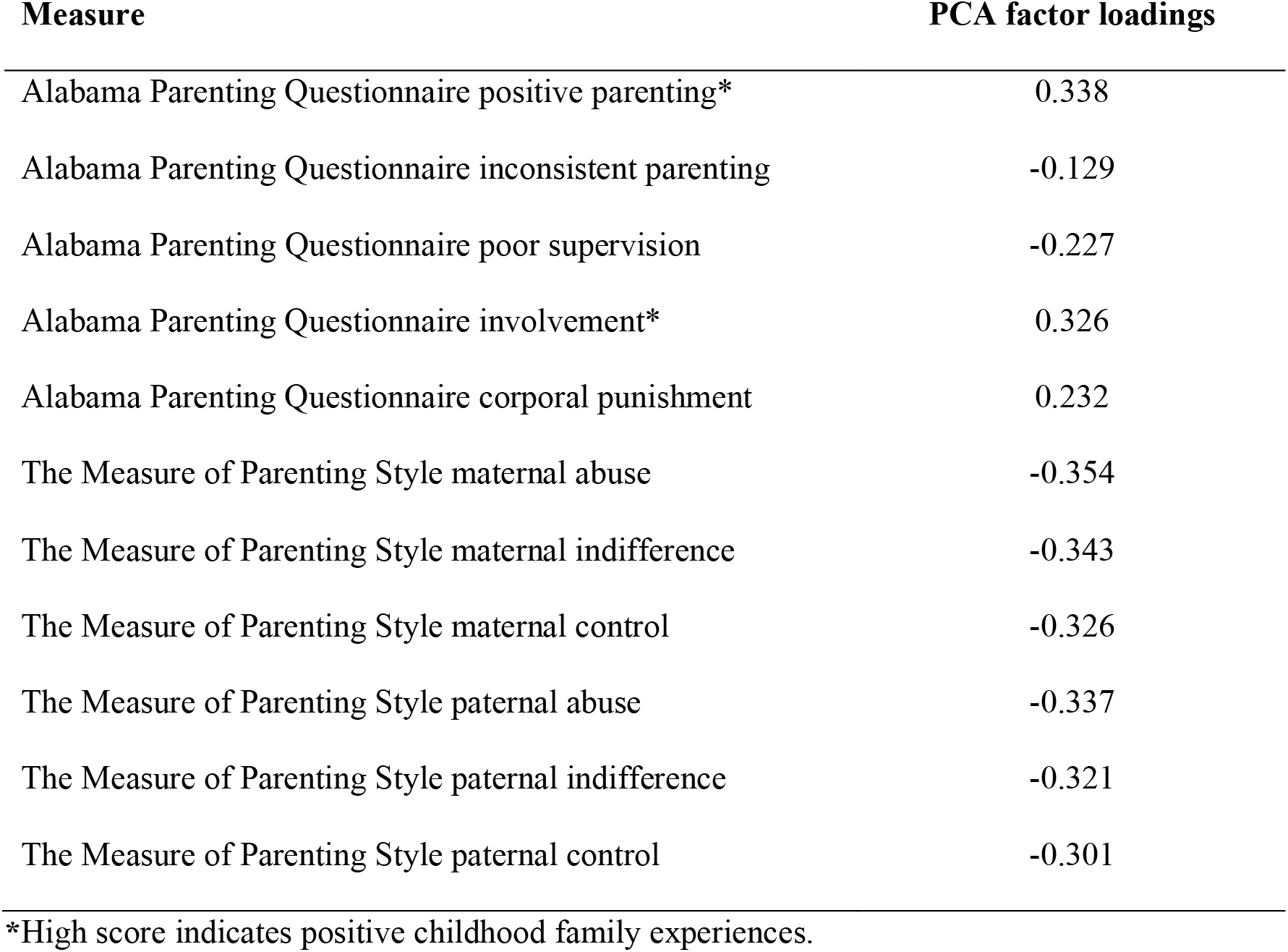
Factor loadings for the PCA’s for childhood family adversity.

| **Measure** | **PCA factor loadings** |
| --- | --- |
| Alabama Parenting Questionnaire positive parenting* | 0.338 |
| Alabama Parenting Questionnaire inconsistent parenting | -0.129 |
| Alabama Parenting Questionnaire poor supervision | -0.227 |
| Alabama Parenting Questionnaire involvement* | 0.326 |
| Alabama Parenting Questionnaire corporal punishment | 0.232 |
| The Measure of Parenting Style maternal abuse | -0.354 |
| The Measure of Parenting Style maternal indifference | -0.343 |
| The Measure of Parenting Style maternal control | -0.326 |
| The Measure of Parenting Style paternal abuse | -0.337 |
| The Measure of Parenting Style paternal indifference | -0.321 |
| The Measure of Parenting Style paternal control | -0.301 |

\*High score indicates positive childhood family experiences.

Other components were observed in the data, however the variance explained by these components was not enough to warrant additional analyses for any of these components in isolation. Scree plots showing the explained variance for each component in both PCA’s can be found in the Supplement (Figure S1). The supplement also includes descriptive statistics for all subscales included in the PCA and demographic variables (age, gender and ethnicity) are listed (Table S3).

### Resilient functioning

To quantify the level of resilient functioning in our sample, we regressed the factor scores for CFA onto the factor score for psychosocial functioning. A linear model provided good fit (adjusted R-squared = 0.28, *F*(1,2404) = 957.2, *p* < 2.2e-16, Est = -06.36e-01, SE = 2.05e-02, *t* = -30.94, *p* < 2e-16, AIC = 10257.64). A quadratic term improved model fit (Est = -0.04, SE = 0.005, *t* = 7.09, *p* = 1.73e-12, AIC = 10204.58), SSM = 204.24, *F*(1) = 50.3, *p* < 1.73e-12). A further cubic model showed weak model fit (Est = -0.002, SE = 0.001, *t* = -1.98, *p* = .05), and only a minimal improvement (AIC = 10202.68, SSM = 15.82 *F*(1) = 3.90, *p* = .05). Therefore, a quadratic model was selected (Figure 1b). Residual scores for this relationship were extracted as they reflect individual degree of resilient functioning and were utilized in the subsequent analyses within the subsample from NSPN that underwent MRI (*N* = 275). These resilient functioning scores were normally distributed in the subsample with MRI data, and there were no significant relationships between estimated resilient functioning and age (*p* = 0.15), gender (*p* = 0.97), SES (*p* = 0.36), or scanning location (*p* = 0.9) (see Supplementary Figure S2).

### Main results

Upon standardization (*z* > 1.65) of those regions that were convergently related with resilient functioning, we found that resilient functioning was associated with a decrease in the nodal degree (*p* < .05 FDR) of the posterior superior temporal sulcus (PSTS), dorsolateral prefrontal cortex (DLPFC), medial prefrontal cortex (MPFC), inferior and middle temporal gyrus (ITG and MTG), lateral occipital cortex (LOCC), pericalcarine, and premotor cortex (Figure 2). Resilient functioning was also related to an increase in the nodal degree (*p* < .05 FDR) of the anterior fusiform gyrus (FG) (Figure 2).

### Sensitivity analysis

To investigate the impact of the statistical correction of regional CT values for age and gender, we next re-estimated structural networks using raw CT values (uncorrected for age and gender, see supplemental material for details). When CT was uncorrected, our findings remained largely similar (Figure S3). Resilient functioning was positively associated with nodal degree in the DMPFC, PSTS, MPFC, left temporal pole, lateral occipital and lingual gyrus and negatively associated with nodal degree in the fusiform gyrus. In addition, our findings now also included positive associations between resilient functioning and nodal degree in the left temporoparietal junction (superior parietal and supramarginal gyrus).

## Discussion

The aim of this study was to investigate how brain structural network topology varies as a function of resilient functioning in a sample of adolescents and young adults with CFA. We showed that resilient psychosocial functioning is negatively associated with nodal degree of the dorsolateral prefrontal cortex (DLPFC), the medial prefrontal cortex (MPFC), the posterior superior temporal sulcus (PSTS), the inferior and middle temporal gyrus (ITG and MTG, resp.), the lateral occipital cortex (LOCC), the pericalcarine cortex, and the premotor cortex. These regions all play a role in a wide array of functions. Of particular interest is the role of these regions in social and emotional processing and regulation, given the importance of socio-emotional functioning in mental health vulnerability in adults exposed to CA (McCrory et al., 2022). The DLPFC is part of the central executive network (CEN) and involved in cognitive control and emotion regulation (Ochsner et al., 2002, 2012). Whereas the MPFC plays an important role in understanding social emotions and mentalizing (Blakemore, 2008; Olson et al., 2013; Van Overwalle, 2009). Importantly, apart from its role in social functioning alterations in medial prefrontal cortex (mPFC)-subcortical circuitry after CFA are associated with a wide array of affective and cognitive functions (Tottenham, 2020). Furthermore, the MPFC and temporal cortex are part of the default mode network (DMN, (Dixon et al., 2017)), which is thought to underpin introspective processes such as emotional processing, decision-making, memory, social cognition, and self-referential processes such as thinking about self-mental states (Northoff et al., 2006; Qin & Northoff, 2011). The DMN is also involved in thinking about other people’s beliefs, intentions, and motivations (Koster-Hale & Saxe, 2013; Spreng et al., 2009). The PSTS has been identified as an important hub in social cognitive processing in different levels, integrating advanced associative and lower-level sensory processing areas (Allison et al., 2000). As such, our findings of reduced nodal degree of regions related to resilient functioning point to regions that help guide socio-emotional functioning.

During adolescence, the brain undergoes remarkable structural and functional reorganization, such as cortical thinning (Frangou et al., 2022; Tamnes et al., 2017; Wierenga et al., 2014), a decrease in cortical surface area and cortical volume (Tamnes et al., 2017) and increases in white matter volume (Giedd et al., 1999). On the microstructural level, such increases in white matter integrity and reorganization of structural brain networks are thought to contribute to more efficient structural brain networks (Koenis et al., 2015). Resting-state functional connectivity shows age-related increases within networks and decreases between networks (Teeuw et al., 2019). In the current sample, decreases in nodal degree have been associated with the pruning of synaptic connections or attenuation of axonal projections in adolescence (Váša et al., 2018) and associated cortical thinning as well as the associated increases in myelination in these regions (Whitaker et al., 2016). As such, decreases in nodal degree are thought to reflect a normative developmental shift to a more efficient brain network configuration (Khundrakpam et al., 2013; Váša et al., 2018). Decreased nodal degree of brain regions in more resilient adolescents thus potentially resembles a mature structural network configuration in these individuals. Social support is known to influence adaptive maturational patterns; stronger mother child interactions are associated with more mature prefrontal-limbic connectivity patterns in children (Gee et al., 2014), and friendship increases are associated with faster cortical thinning in the mPFC in adolescents (Becht et al., 2021). Friendships support has been well established as predictor of resilient functioning in adolescent and young adults exposed to CA (Fritz et al., 2018; van Harmelen et al., 2017; 2020). Taking together the evidence of previous studies in the same and different samples, we suggest that our findings of negative associations between nodal degree and resilient functioning may reflect a more mature-like structural network topology in more resilient individuals. However, as we did not include longitudinal data, we can only speculate that developmental mechanisms explain the found associations. As such, it should be examined whether lower nodal degree in individuals with more resilient functioning reflect more mature structural brain topology and or distinct developmental trajectories of the DLPFC, MPFC, PSTS, ITG and MTG.

Resilient functioning was also associated with increased nodal degree in the anterior fusiform gyrus. One interpretation for our finding would be that maturation of the fusiform gyrus might be protracted (Haist & Anzures, 2017) or that this structure may be less mature in resilient individuals at this age group. The fusiform gyrus is specialized in the processing of faces and involved in facial emotion recognition (Adolphs, 2002). Aberrant neural activation in response to emotional faces in individuals exposed to CA is consistently reported in literature (Bérubé et al., 2023). The social transactional model of psychiatric vulnerability in adults exposed to CA suggests that such aberrant facial emotion processing, reflects biased threat processing which could inadvertently impact social functioning and relations, and thereby increase vulnerability for psychiatric disorders (McCrory et al., 2022). Indeed, functional imaging studies showed that resilient adults with CA show improved ability to regulate emotions through medial prefrontal cortex–limbic downregulation, lower hippocampal activation to emotional faces, and increased amygdala habituation to stress (reviewed in Moreno-Lopez et al., 2020). It should be examined if increased nodal degree in the anterior fusiform gyrus aids resilient functioning through improved facial emotion processing.

Strengths of this study include a large sample of carefully assessed participants recruited from the community with low to moderate CFA experiences and the use of a multi-parameter mapping protocol. One limitation is that we focused on CFA limiting the generalizability of results to individuals exposed to other types of CA. Furthermore, individuals with current treatment for psychiatric disorders were excluded and the lower spectrum of CFA was overrepresented in this study. Therefore, future studies are needed to investigate whether similar or distinct mechanisms aid resilient functioning in individuals with more extreme CFA experiences. A limitation of our residual variance approach to quantify resilient functioning is that this entails a strong association between psychosocial functioning and the measures of functioning; as the residuals will, by design, be highly correlated with psychosocial outcomes. However, in our sample CFA severity was correlated significantly with psychosocial outcomes as such, our approach can explicitly separate functioning outcomes towards the extremes of CFA severity. As an example, an individual who has experienced little or no CFA will have lower resilient functioning scores than an individual who experienced severe CFA, even if the latter may have lower absolute psychosocial functioning (see Figure 1b). Another limitation of our study is that we did not include subcortical regions in our analysis. Future studies should aim to explore structural covariance between and among cortical and subcortical grey matter structures to describe a more substantive and thorough structural covariance network underlying possible differences in for instance social cognition and emotional regulation in adolescents. Also, it was beyond the scope of this study to incorporate different structural covariance measures while other measures, such as modularity, show interesting developmental effects as well (e.g., Aboud et al., 2019). Further, the results of any sliding window method are dependent on the parameters used (i.e., window width and step size), in addition to the resilience scores being non-homogeneously distributed. However, we systematically varied these parameters, and focused on only those regions showing significant convergent results across ten parameter combinations.

CA is one of the strongest predictors of mental health problems in later life, and as such it is critical that we better understand how resilience can be achieved in young people with CA. To do so, resilience research examines why some young people with CA go on to develop mental illness, whereas others do not. By better understanding this variability in functioning outcomes in CA-exposed individuals, resilience research represents a shift away from a disease focussed model towards a preventative model. As such, resilience research aims to inform intervention and prevention efforts for individuals at risk (Masten et al., 2019; Luthar & Cicchetti, 2000). Recent models emphasise that resilience is facilitated by complex interrelations across cultural, social, psychological, and neurobiological systems and their development over time (Masten & Cichetti, 2010; Masten et al., 2021). Within this multisystem framework, neuroimaging studies help inform our understanding of the neurobiological mechanisms that help aid resilient functioning (Ioannidis et al., 2020). By studying how these mechanisms interact with social or psychological systems these studies can provide key insights for prevention or intervention efforts (Cicchetti & Toth, 2015). In this study, we integrated two systems: individual psychosocial functioning and the brain. We examined the structural network topology of resilient functioning in adolescents and young adults exposed to CFA. We found that higher resilient functioning was convergently associated with lower nodal degree of DLPFC, MPFC, PSTS, ITG and MTG, LOCC, pericalcarine cortex, and the premotor cortex. These regions all play a role in a wide array of functions, of particular interest is their role in social-emotional functioning. Developmental changes in adolescence include extensive remodelling of structural covariance patterns, including reductions in nodal degree. As previous work in this sample showed negative associations between age and nodal degree (Váša et al., 2018; Whitaker et al., 2016), our findings of lower nodal degree being related to higher resilient functioning may be compatible with more mature-like structural network topology in more resilient young people with CFA.

## Acknowledgments and funding sources

The authors are grateful to the volunteers who took part in these studies, as well as all the members of staff involved in the recruitment process. This work was supported by the Neuroscience in Psychiatry Network (NSPN) Consortium, a strategic award from the Wellcome Trust to the University of Cambridge and University College London (095844/Z/11/Z); the Leiden Social Resilience and Security programme, the Cambridge NIHR Biomedical Research Centre and by the Max Planck–UCL Centre for Computational Psychiatry and Ageing, a joint initiative of the Max Planck Society and University College London; a Royal Society Dorothy Hodgkin Fellowship for Prof. Anne-Laura van Harmelen (DH150176); an NIHR Senior Investigator award for Prof. Ed Bullmore and a Wolfe Health Fellowship for Dr. Laura Moreno-López. Dr. František Váša was supported by the Gates Cambridge Trusts, the Data to Early Diagnosis and Precision Medicine Industrial Strategy Challenge Fund, UK Research and Innovation (UKRI) and the Bill & Melinda Gates Foundation.

## Appendix – Supplementary methods and results

### 1. Supplementary methods

#### 1.1. Measures used to calculate resilient psychosocial functioning

The Mood and Feelings questionnaire (MFQ; 66) is a 33 item self-report questionnaire measuring depressive symptoms within the past two weeks. Responses are rated on a 3-point Likert scale (i.e., “Not true,” “Sometimes,” and “True”). Higher MFQ sum scores indicate more severe depressive symptomology. At baseline, internal consistency of the MFQ is excellent (Cronbach’s alpha = 0.91 - 0.93).

The Revised Children’s Manifest Anxiety Scale (RCMAS; 67) is a 49 item self-report questionnaire which assesses global anxiety with five subscales: physiological anxiety, worry, social anxiety, defensiveness, and inconsistent responding index. Responses range on a 4-point Likert scale from “Never” to “Always,” with higher sum scores indicating higher levels of anxiety. Internal consistency of the RCMAS was excellent at baseline (Cronbach’s alpha = 0.94).

The Short Leyton Obsessional Inventory (S-LOI; 68) is an 11-item questionnaire which assesses obsessional/anxious symptomology indicative of adolescent obsessive-compulsive disorder (OCD). The S-LOI comprises three subscales: compulsions, obsessions/incompleteness, and cleanliness. Responses range from “Never” to “Always” on a 4-point Likert scale. Higher sum scores denote higher levels of obsessional symptoms. Internal consistency of the S-LOI is good (Cronbach’s alpha = 0.84).

The Child Behavior Checklist (CBCL; 69) is an 11 item self-report questionnaire which screens for symptoms of antisocial behavior conceptually grounded in DSM-IV conduct disorder (CD) items. Responses were on a 4-point Likert scale from “Never” to “Always.” In our analysis, endorsement of the two highest responses in agreement were combined. At baseline, the internal consistency of the CBCL was good (Cronbach’s alpha = 0.74).

The Kessler Psychological Distress scale (K10; 70) is a 10-item questionnaire which yields a global measure of distress encompassing both anxious to depressive symptomology. Responses range on a 5-point Likert scale from “None of the time” to “All of the time.” The higher one scores on the K10, the more psychological distress is indicated. Internal consistency at baseline of the K10 was high (Cronbach’s alpha = 0.89).

The Antisocial Process Screening Device (APSD; 71) is a 20-item scale measuring psychopathic personality traits. Possible responses on the ASPD are “Not at all true,” “Somewhat true,” and “Certainly true”; with higher sums scores on the ASPD indicating higher levels of psychopathy. Internal consistency of the ASPD is good (Cronbach’s alpha = 0.73).

The Child and Adolescent Dispositions Scale (CADS-Y; 72) measures three dispositional traits: sympathy for others, negative emotional responsivity, and positive reactivity to risk and novelty which may predispose one to CD. Participants were asked to rate how closely they are described by an item, with responses ranging on a 4-point Likert scale from “Not at all” to “Very much/Very often.” Internal consistency at baseline for sum scores of the three dimensions was good (alphas = 0.78, 0.72, 0.77, respectively).

The Inventory of Callous-Unemotional Traits (ICU; 73) is a 24-item scale which comprises three trait subscales (i.e., callousness, uncaring, unemotional) intended to measure indications of psychopathy. Participants are asked to rate of a 4-point Likert scale ranging from “Never/Almost Never” to “Always/Almost Always” how closely the statements related to these personality traits described them. Higher scores indicated higher levels of psychopathy. Internal consistency of the ICU is good (Cronbach’s alpha = 0.82).

The Schizotypal Personality Questionnaire (SPQ; 74) is a 74-item scale which measures nine subscales of schizotypy (i.e., ideas of reference, excessive social anxiety, odd beliefs or magical thinking, unusual perceptual experiences, odd or eccentric behaviour, no close friends, odd speech, constricted affect, and suspiciousness) in non-clinical samples. Response choices are dichotomized “Yes”/“No.” A higher sum score on the SPQ indicates more schizotypal symptomology. The SPQ has high internal consistency (Cronbach’s alpha = 0.91).

The Barratt Impulsivity Scale (BIS-11; 75) is a 30-item scale measuring the behavioural/personality components of impulsivity (i.e., attentional, motor, and non-planning). With response choices ranging from “Rarely/Never” to “Always,” participants were requested to select the a choice which most closely resembles how they behave. Higher scores on the BIS suggest higher levels of impulsiveness. Internal consistency for the BIS is generally good (Cronbach’s alpha range= 0.79 - 0.83).

The Warwick-Edinburgh Mental Well Being Scale (WEMWBS; 76) has 14 items used to address mental wellbeing. Participants were to answer how accurately each statement described their experiences within the last two weeks. On a 5-point Likert scale, responses ranged from “none of the above” to “all of the above.” Here, higher scores on the WEMWBS indicate greater mental well-being. Internal consistency for the WEMWBS is excellent at baseline (population sample Cronbach’s alpha = 0.91).

#### 1.2. Measures used to calculate childhood family adversity

The Measure of Parenting Style (MOPS; 77) is a 12 item self-report measure that assesses perceived parenting styles across three domains; abuse, indifference, over-control. Participants were asked to rate both their mother’s and father’s parenting behaviour on 15 statements, on a 4-point scale. The full response range is “not true at all”, “slightly true”, “moderately true”, “extremely true”. The ‘abuse’ scale consisted of 5 items, asking whether maternal/paternal behaviours were verbally abusive, unpredictable, physically violent, elicited feelings of danger, or elicited feelings of lack of safety. The ‘overly controlling’ scale consisted of 4 items where maternal/paternal behaviour was overprotective, over controlling, critical, or made the participant feel guilty. Finally, the ‘indifference’ scale assessed 6 items of maternal/paternal behaviour where the parent was ‘ignoring, uncaring, rejecting, uninterested in, would forget about, or would leave the participant on his/her often. Sum scores to responses in these items were calculated with higher scores representing more abusive, over controlling, or indifferent behaviour reported. Internal consistency was good for the maternal subscales (Cronbach’s alpha maternal over control = 0.70, indifference = 0.86, abuse = 0.78). For paternal parenting, the internal consistency at baseline ranged from acceptable (Cronbach’s alpha paternal over control = 0.65) to excellent (Cronbach’s alphas paternal abuse = 0.88, paternal indifference = 0.93).

The Alabama Parenting Questionnaire (APQ; 78) measures parenting practices. We used the 9-item short-form and added the ‘Corporal Punishment’ (3 items) and ‘Involvement’ scale (3 items). Participants were asked to rate how typical each item occurred or used to occur in their family home on a five-point scale ranging from “never”, “almost never”, “sometimes”, “often” to “always”. We calculated sum scores for the five subscales: Corporal Punishment, Positive Parenting, Inconsistent Discipline, Poor Supervision, and Involvement, with higher scores reflecting higher frequency of the behaviour. Thus, high scores can indicate positive parenting (i.e., involvement, positive parenting), or negative parenting (i.e., inconsistent discipline, poor supervision, corporal punishment). Internal consistency at baseline was acceptable (Inconsistent discipline & poor supervision: Cronbach’s alpha’s > 0.62), and good (Positive parenting, Involvement, Corporal Punishment Cronbach’s Alpha’s > 0.71).

#### 1.3. Regional changes in unadjusted cortical thickness as a function of resilience

To investigate the impact of the statistical correction of regional CT values for age and gender, we re-estimated structural networks using raw CT values (uncorrected for the potentially confounding variables above). All other steps involved in structural network construction were consistent with the procedure described in the main text. Briefly, we used Pearson correlations on raw CT values (uncorrected for age and gender) to construct structural networks for overlapping subsets of participants (“windows”) ordered by increasing resilience. Next, we used bootstrapping to threshold the networks. Using this method, and for each window, an equal number of participants were resampled with replacement to construct 1000 bootstrapped structural networks. We then examined whether there were significant relations between each pair of regions across all bootstrapped networks. Consistent relationships between a pair of regions (at *p* < .001 FDR-adjusted at the pair level) were retained and the remaining relationships were discarded. We then assessed network topology focusing on degree only.

### 2. Supplementary Results

#### 2.1. Supplementary tables

**Table S1.** Correlation Resilient functioning scores from the 5 imputed datasets.

| Imputed datasets |  |  |  |  |
| --- | --- | --- | --- | --- |
|  | 1 | 2 | 3 | 4 |
| 2 | $r=1, t(2404)=23267, p<2.2e-16$ | na | | |
| 3 | $r=99, t(2404)=440.83, p<2.2e-16$ | $r=99, t(2404)=440.83, p<2.2e-16$ | na | |
| 4 | $r=99, t(2404)=497.6, p<2.2e-16$ | $r=99, t(2404)=497.6, p<2.2e-16$ | $r=99, t(2404)=431.86, p<2.2e-16$ | na |
| 5 | $r=99, t(2404)=456.42, p<2.2e-16$ | $r=99, t(2404)=456.42, p<2.2e-16$ | $r=99, t(2404)=416.73, p<2.2e-16$ | $r=99, t(2404)=448.14, p<2.2e-16$ |

**Table S2.** Raw cortical thickness correlated with resilient functioning (uncorrected).

| Region | X | Y | Z | $\beta$ |
| --- | --- | --- | --- | --- |
| Left banks of the superior temporal sulcus | -53 | -50 | 8 | -2.5 |
| Left lingual | -7 | -77 | -1.6 | -2 |
| Right medial orbitofrontal | -7 | 53 | -11 | -2.5 |
| Left parahippocampal | -22 | -25 | -25 | -3 |
| Left paracentral | -9 | -21 | 49 | -5 |
| Left postcentral | -16 | -36 | 71 | -2 |
| Left superior parietal | -32 | -47 | 59 | -2 |
| Left superior parietal | -21 | -61 | 62 | -3 |
| Left supramarginal | -54 | -47 | 39 | -3 |
| Left supramarginal | -55 | -33 | -37 | -3.5 |
| Left temporal pole | -32 | -11 | -35 | -3.6 |
| Right fusiform | 35 | -8 | -36 | 2 |
| Right inferior temporal | 52 | -33 | -25 | -3 |
| Right lateral occipital | 43 | -79 | -11 | -2 |
| Right lateral occipital | 29 | -87 | -12 | -3 |
| Right lateral occipital | 42 | -84 | -1 | -3 |
| Right paracentral | 5 | -17 | 61 | -1 |
| Right pericalcarine | 15 | -71 | 8 | -2 |
| Right precentral | 54 | 6 | 24 | -4.8 |
| Right precentral | 23 | -16 | 66 | -3.4 |
| Right rostral middle frontal | 32 | 30 | 38 | -2.3 |
| Right superior frontal | 19 | 3 | 60 | -3.2 |

**Table S3.**
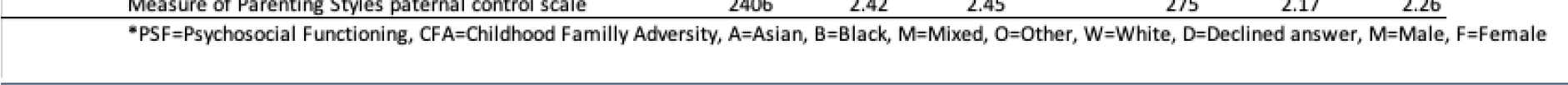
Socio-demographics and descriptives of psychosocial functioning and childhood family adversity subscales.

#### 2.2. Supplementary figures

**Figure S1.**
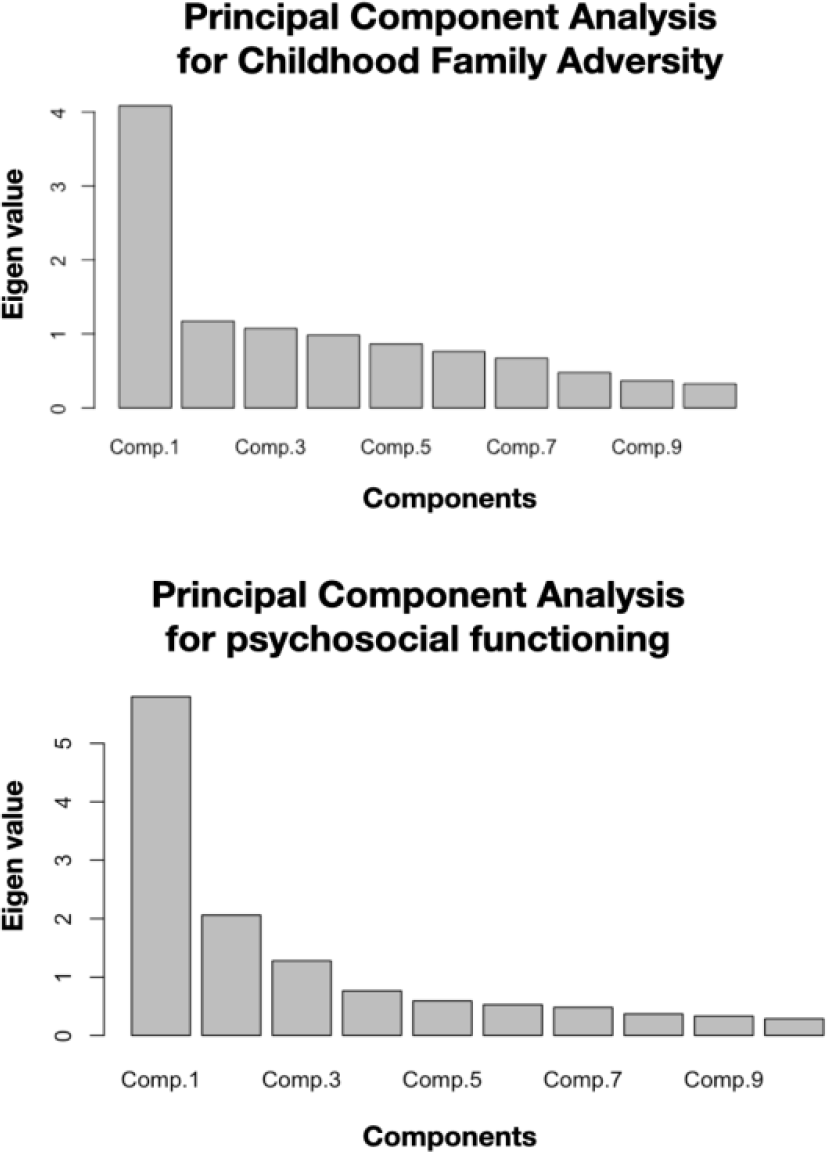
Scree plots of eigen values for each component in both PCA’s.

**Figure S2.**
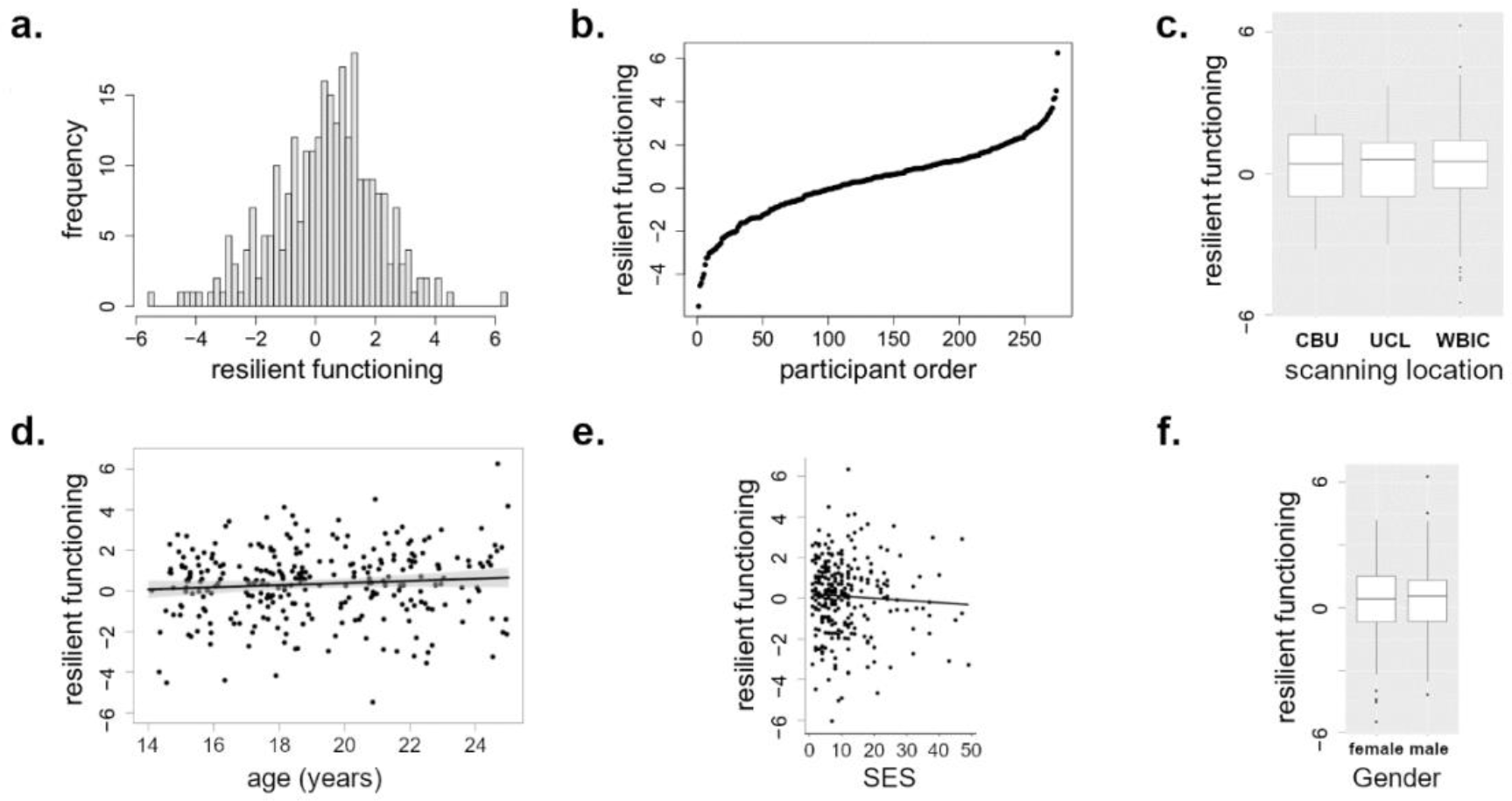
Homogeneity in resilient functioning-distribution of the participants. A) Distribution of resilient functioning in the participants; B) Resilient functioning-gaps between consecutive participants; C) Association between resilient functioning and scanning location. D) Correlation between resilient functioning and age; E) Association between resilient functioning and socioeconomic status; F) Association between resilient functioning and gender. CBU=Cognition and brain science unit; UCL=University college London; WBIC=Wolfson brain imaging center; SES=Socioeconomic status.

**Figure S3.**
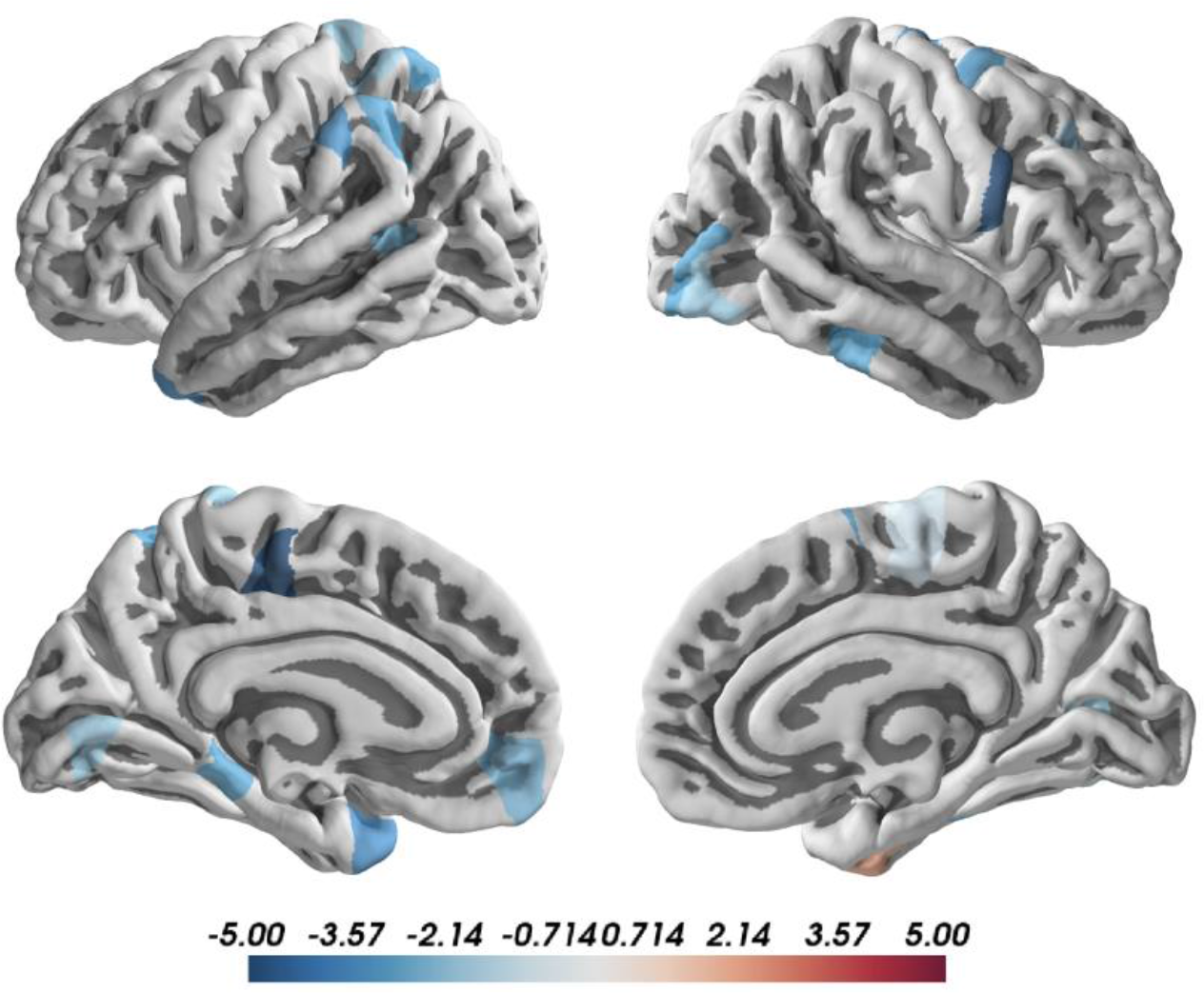
Convergence indices for the relation between resilient functioning and nodal degree based on cortical thickness measures uncorrected for age and gender. Positive (red) and negative (blue) correlations between resilient functioning and node degree cortical thickness measures uncorrected for age and gender.

